# Effect of Perfluorooctanesulfonic acid (PFOS) on immune cell development and function in mice

**DOI:** 10.1101/2020.01.20.913293

**Authors:** Luisa Torres, Amie Redko, Candice Limper, Brian Imbiakha, Sue Chang, Avery August

**Affiliations:** Department of Microbiology & Immunology, Cornell University, Ithaca, NY 14853, USA; 3M Company, St. Paul, MN 55144, USA

**Keywords:** PFOS, influenza, immune system

## Abstract

Perfluorinated compounds, such as Perfluoroctane sulfonic acid (PFOS) are compounds containing carbon chains where hydrogens have been replaced with fluorines, and belong to a larger family known as Per- and polyfluoroalkyl substances (PFAS). The strength of the carbon–fluorine bond makes perfluorinated compounds extremely resistant to environmental degradation. Due to the persistent nature of PFOS, research has been directed to elucidating possible health effects of PFOS on humans and laboratory animals. Here we have explored the effects of PFOS exposure on immune development and function in mice. We exposed adult wild-type mice to 3 and 1.5 μg/kg/day of PFOS for 2 and 4 weeks, respectively, and examined the effects of PFOS exposure on populations of T cells, B cells, and granulocytes. These doses of PFOS resulted in serum levels of approximately 100 ng/ml with no weight loss during exposure. We find that PFOS does not affect T-cell development during this time. However, while PFOS exposure reduced immune cell populations in some organs, it also led to an increase in the numbers of cells in others, suggesting possible relocalization of cells. We also examined the effect of PFOS on the response to influenza virus infection. We find that exposure to PFOS at 1.5 μg/kg/day of PFOS for 4 weeks does not affect weight loss or survival, nor is viral clearance affected. Analysis of antibody and T cell specific antiviral responses indicate that at this concentration, PFOS does not suppress the immune cell development or antigen specific immune response.

## Introduction

Perfluorooctanesulfonyl fluoride (POSF)-based compounds are a class of surfactant chemicals that have been used in a wide variety of industrial and consumer applications^1^. Perfluorooctanesulfonate (PFOS, C^8^F^17^SO^3-^) is the end-stage metabolite for POSF-based chemistry and the strength of the carbon–fluorine bond makes it extremely stable and resistant to environmental degradation^2^. PFOS is ubiquitously distributed in the environment and does not metabolize under normal physiological conditions, with bioaccumulation potential in both wildlife and humans. Observations in multiple species indicate that PFOS is slowly eliminated with serum half-lives of approximately 1 – 2 months for rodents, 4 months for non-human primates, and 3 – 5 years for humans^3–5^.

Due to the persistent nature of PFOS, research has been directed at evaluating possible health effects of exposure to PFOS^6^. In epidemiological studies in humans, exposure to PFOS have been reported to be associated with several health effects, and meta-analyses and/or critical reviews of this literature suggest association with lower birthweight^7 8^, infertility^9^, altered thyroid function in pregnant women and children^10^; higher cholesterol^11^, and immunological effects^12^. With regards to potential effects on the immune system, a systematic review of the literature was conducted by the National Toxicology Program (NTP), which concluded that PFOS is “presumed to be an immune hazard to humans”^12^ based on reports of PFOS-mediated suppression of the antibody response in animal studies, and epidemiological studies in humans. While the evidence for an effect of PFOS on the immune system was based on findings of suppression of the antibody response and was found to be substantial by the NTP, there was less evidence from experimental animals that PFOS suppresses disease resistance and natural killer (NK) cell activity. Furthermore, the NTP reports indicate that the mechanism(s) for such PFOS-associated immunotoxicity was not well-understood. In addition, among the supporting studies cited by NTP, the seemingly high level of PFOS dose(s) used in the studies with laboratory animals often resulted in significant body weight loss^12^, and the accompanying stress and elevated levels of corticosteroids (a known confounder for immune response^13^), that accompanies these high dose treatments cannot be excluded^12^.

Here we have evaluated the potential immunotoxicity of exposure to PFOS in wild-type C57Bl/6 mice after repeated oral treatments for either 2 weeks or 4 weeks, using doses of PFOS that did not lead to weight loss in the animals. We examined the development of immune cells, including T cell development in the thymus, as well as B cell and granulocyte development in the bone marrow and spleen. We also evaluated the effect of PFOS exposure on the immune response to influenza virus infection.

## Materials and Methods

### Material

All chemicals used in this study were reagent-grade. Potassium perfluorooctanesulfonate (K^+^PFOS) was obtained from 3M Company (St. Paul, MN). The details of its purity (88.9%) and analytical characterization are described in Chang et al.^14^.

### PFOS dose formulation and analysis

K^+^PFOS was prepared in vehicle (0.5% Tween 20 in deionized water) at either 0.3 μg/mL or 0.6 μg/mL; and with a daily dosing volume of 5 mL per kg body weight, it corresponded to a daily targeted dose of either 1.5 μg/kg body weight/day or 3 μg/kg body weight /day, respectively. The dosing solutions and the serum samples were analyzed for PFOS concentration by LC-MS/MS based on previous methods^14, 15^. Briefly, matrix-matched standard curves were prepared and ^18^O_2_-PFOS was used as internal standard for all extractions. All standards and samples underwent solid phase extraction prior to LC-MS/MS analyses. API 6500 mass spectrometer (AB SCIEX, Redwood City, California) with Turbo Ion Spray (pneumatically assisted electrospray ionization source) that operated in a negative ion mode was used for this study, and chromatography analysis was conducted with a Mac-Mod ACE C-18 HPLC column (2.1 mm *i.d*. × 75 mm, 5 µm particle size) at 0.25 mL/min under gradient condition (95%/5% 2 mM ammonium acetate/acetonitrile and acetonitrile). All source parameters were optimized under the conditions according to manufacturer’s guidelines. Transition ions monitored were: (1) PFOS: 499 → 80 m/z; and (2) Internal Standard ^18^O_2_-PFOS: 503 → 84 m/z. Concentration of PFOS in the vehicle solution (0.5% Tween 20 in water) was also measured and was below the lower limit of quantification of 5 ng/mL.

### Mice

Adult male and female wild-type C57BL/6 mice bred at Cornell University were used this study. The mice were approximately eight weeks old and weighed approximately 20-25 grams at the start of the treatment. All experimental procedures were reviewed and approved by the Institutional Animal Care and Use Committee and Institutional Review Board at Cornell University and conformed to Guide for the Care and Use of Laboratory Animals^16^.

### Study design for PFOS exposure

#### 2-week PFOS exposure

Male and female C57Bl/6 mice (n=6-7/group) were administered daily with either vehicle (0.5% Tween 20 in water) or K^+^PFOS (referred to hereafter as PFOS) at 3 μg/kg/day for 14 days via oral gavage. The theoretical total administered dose (TAD) was 42 μg PFOS/kg body weight after 14 daily oral administrations. At the end of the exposure period (one day after last dose), animals were euthanized under CO_2_ asphyxiation and blood was collected from the abdominal aorta. Blood samples were left at room temperature for 40 minutes and processed to serum after centrifugation (2500 rpm, 20 min). Serum samples were stored frozen at -80°C pending further analysis. Thymus, spleen, liver, Peyer’s patches, and bone marrow were also collected from all mice and further processed for lymphocyte isolation as described below (see organ collection and lymphocyte isolation).

#### 4-week PFOS exposure

Male and female C57Bl/6 mice (n=4-7/group) were administered daily with either vehicle (0.5% Tween 20 in water) or PFOS at 1.5 μg/kg/day for 28 days via oral gavage. After 28 consecutive daily treatments, the theoretical total administered dose (TAD) was 42 μg PFOS/kg body weight as well. At the end of the exposure period (one day after last dose), animals were euthanized and blood and organs collected as previously described for 2-week PFOS exposure.

#### Influenza virus infection post 4-week PFOS treatment

Both male and female C57Bl/6 mice were used in this study. In experiment #1, mice (n=4-5/group) were administered daily with either vehicle (0.5% Tween 20 in water) or PFOS at 1.5 μg/kg/day for 28 days via oral gavage. One day after the last vehicle or PFOS treatment, mice were infected with mouse influenza virus strain A/WSN/33(H1N1) intranasally (1e6 pfu/mouse). Body weights were monitored daily for 11 days post infection followed by euthanization. Animals were euthanized as previously described and bronchoalveolar lavage collected from lungs, and blood, lungs, liver and spleen collected as described above. Serum and BAL samples were stored frozen at -80°C until analyzed.

In experiment #2, mice (n=10/group) were administered daily with either vehicle (0.5% Tween 20 in water) or PFOS at 1.5 μg/kg/day for 28 days via oral gavage. One day after the last vehicle or PFOS treatment, mice were infected with mouse influenza virus strains A/WSN/33(H1N1)(WSN-OVA_I_) and influenza A/WSN/33(H1N1)(WSN-OVA_II_) intranasally (2e4 pfu/mouse). Influenza A/WSN/33(H1N1)(WSN-OVA_I_) is a recombinant influenza virus carrying the peptide epitope OVA_257–264_ from the model antigen Ovalbumin, and influenza A/WSN/33(H1N1) (WSN-OVA_II_) was a recombinant influenza virus carrying the peptide epitope OVA_323–339_ from the model antigen Ovalbumin^17, 18^. Body weights, euthanization and tissue collections were as described above.

### Organ collection and lymphocyte isolation

Thymus, spleen, liver, Peyer’s patches, and bone marrow were collected from euthanized mice. The spleen and thymus where passed through a 100-μm cell strainer and crushed with the plunger of a 3 ml syringe to get a single-cell suspension that was collected in 5 ml of complete RPMI medium. The liver was injected with 10 ml of PBS followed by 5 ml of 0.05% collagenase A and Dispase II in complete medium. The homogenized tissue was filtered and centrifuged for 5 min at 400 rcf. 5 ml of Lympholyte-M was added to the 5 ml of PBS containing the resuspended pellet and centrifuged for 20 minutes at 1200 rcf. Lymphocytes were obtained from the interface layer. Lymphocytes were isolated from the Peyer’s patches as previously described^19^. The bone marrow was flushed with 1 ml of complete RPMI medium using a 1 ml syringe and a 23-gauge needle to obtain a single cell suspension of lymphocytes. Cells were counted with an automated cell counter, surfaced-stained, and analyzed by flow cytometry.

### Hemagglutination inhibition (HAI) assays

Hemagglutination inhibition assays were performed as described ^20^. In brief, a 1% (v/v) suspension of chicken red blood cells in PBS was used to assay hemagglutination of influenza virus and 2 fold dilutions of serum from mice infected with the mixture of WSN-OVA_I_ and WSN-OVA_II_. Sera were incubated with 4 hemagglutination units of WSN virus. The HI titer was defined as the reciprocal of the highest serum dilution inhibiting hemagglutination.

### Antibodies for flow cytometry

T- cell panel: APC-Cy7- TCRβ (BD Biosciences), eFluor 450-CD4, PerCP-Cy5.5-CD8α, PE NKT, PeCy7 NK1.1, FITC CD44, AF700 CD62L, APC TCRγδ. Granulocyte panel: Pacific Blue Ly6G, PE Siglec-F, APC CD11c, AF700 MHCII, CF594 CD11b, APC Cy7 c-kit, BV510 F4/80, PeCy7 FcεR1*α*. B cell panel: AF700 B220, APC Cy7 IL17R, PerCP-Cy 5.5 CD24, PE CD21, PECy7 CD23, APC IgM. iNKT cells were identified using CD1d/*α*-galactosyl ceramide tetramers from the NIH Tetramer Core Facility (https://tetramer.yerkes.emory.edu/). OVA-antigen specific T cells in influenza A/WSN/33(H1N1) (WSN)-OVA_I_ influenza A/WSN/33(H1N1) (WSN)-OVA_II_ infected mice were identified using MHC-class II and class I OVA/tetramers that identify OVA-specific CD4^+^ and CD8^+^ T cells^21 22^, also obtained from the NIH Tetramer Core Facility (https://tetramer.yerkes.emory.edu/).

### Flow Cytometry

Surface staining of live cells was done in the presence of anti-CD16/32 (clone 93; Fc block) and fixable viability dye. Flow cytometry data were acquired on LSRII, FACS Aria II or Fusion systems (BD Biosciences), and analyzed in FlowJo (Tree Star, Ashland, OR). All analyses were performed on fixable viability dye negative singlet population.

### Statistical analysis

All statistics were performed using GraphPad Prism 8 for Mac. ANOVA or two-tailed Student’s *t* tests with Welch’s correction for samples having possibly unequal variances was used to determine significance between control and experimental groups. Data were presented as mean ± SEM. Data were considered statistically significant when p<0.05.

## Results

### PFOS concentrations in serum samples after 2-week or 4-week PFOS treatments

Male and female mice were orally administered with either vehicle or PFOS daily for 14 days (3 μg/kg/day) or 28 days (1.5 μg/kg/day), in order to achieve comparable serum PFOS levels. After either 14 days or 28 days of PFOS exposure at 3 μg/kg/day or 1.5 μg/kg/day, respectively, the mean serum PFOS concentrations were (116 ng/m; ± 2.8 (SEM)) ng/mL and (99.6 ± 4.4 (SEM)) ng/mL, respectively There was a statistical significant difference between the mean serum PFOS levels of the 2 and 4 week exposures (**Fig. 1A**).

**Figure 1.**
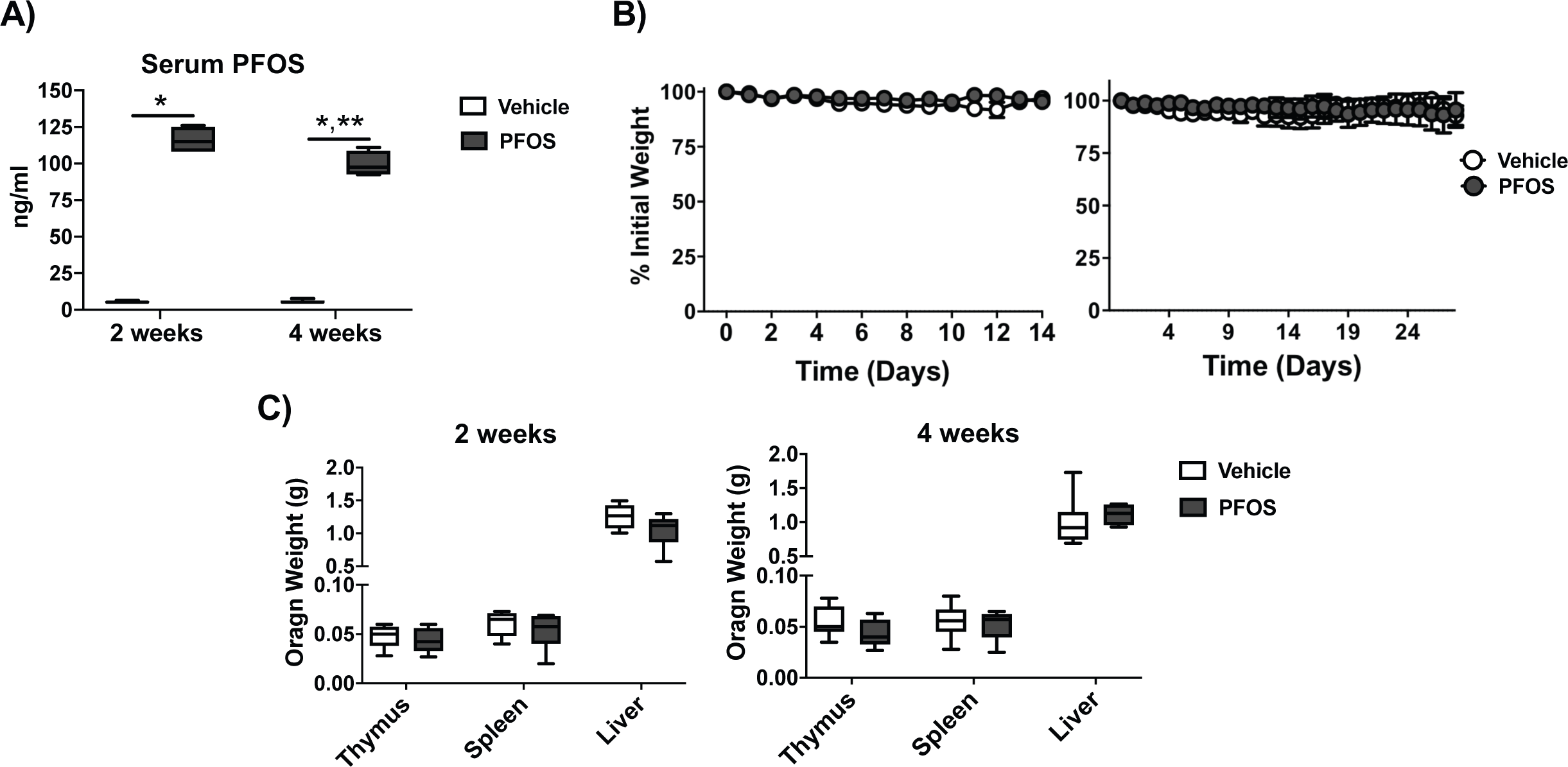
PFOS exposure at 3 μg/kg/day PFOS for 2 weeks or 1.5 μg/kg/day for 4 weeks does not significantly affect body weight. (**A**) WT C57Bl/6 mice were exposed to vehicle or PFOS at 3 μg/kg/day for 2 weeks or 1.5 μg/kg/day for 4 weeks by oral gavage. At the end of the exposures, serum PFOS concentrations were determined. Mean ± SEM, n=6 (vehicle exposed) and n=7 (PFOS exposed) for 2 weeks of exposure, *p<0.0001 vs vehicle 2 weeks exposed mice, Mean ± SEM, n=7 (vehicle exposed) and n=4 (PFOS exposed), *p<0.0001 vs vehicle 4 weeks exposed mice, *p<0.0005 2 week vs 4 week PFOS exposed mice. (**B**) Mice in (A) were weighed daily and percent original weight plotted. 2 weeks (left panel) or 4 weeks (right panel), mean ± SEM, n=7 (vehicle exposed) and n=8 (PFOS exposed) for 2 weeks of exposure and n=7 (vehicle exposed) and n=5 (PFOS exposed) for 4 weeks of exposure. (**C**) The indicated organs from the mice described in (A) were weighed. Mean ± SEM, n=5 (vehicle exposed) and n=6 (PFOS exposed) for 2 weeks of exposure and n=7 (vehicle exposed) and n=5 (PFOS exposed).

### PFOS exposure at 3 μg/kg/day PFOS for 2 weeks or 1.5 μg/kg/day for 4 weeks does not affect body weight or selected organ weights

The body weight of the PFOS-exposed mice were monitored daily. After 2 weeks or 4 weeks of PFOS exposure at 3 μg/kg/day or 1.5 μg/kg/day, respectively, we did not observe significant changes in body weight at the end of either treatment period (**Fig. 1B)**. Analysis of the thymus, spleen, and liver indicated that at these concentrations, PFOS does not affect the weights of these organs (**Fig. 1C**).

### Effect of 2-week or 4-week exposure to PFOS on T cell development in the thymus

T lymphocytes develop from hematopoietic stem cells that differentiate into lymphoid progenitors in the bone marrow, which migrate to the thymus, where they initially lack most of the surface molecules characteristic of mature T cells. These cells give rise to populations of αβ T cells and γδ T cells. T cells are initially double negative for co-receptors CD4 and CD8 and later differentiate into αβ T cells and γδ T cells, with a CD4/CD8 double positive intermediate stage before becoming single positive CD4 and CD8 αβ T cell depending on the class of MHC molecules they encounter, followed by exit from the thymus and circulation in the periphery^23^. To determine the effect of PFOS exposure on T lymphocyte development, we analyzed thymocytes in the C57BL/6 male and female mice exposed to 3 μg/kg/day PFOS for 2 weeks or 1.5 μg/kg/day for 4 weeks. Regardless of exposure duration, PFOS did not affect the numbers of thymocytes (**Fig. 2A**). Flow cytometric analysis of thymocyte subpopulations (**Fig. 2B**), indicated that exposure also did not affect the percentages of double negative or double positive thymocytes for CD4 and CD8, indicating that PFOS exposure at this concentration did not affect T cell development (**Fig. 2C**).

**Figure 2.**
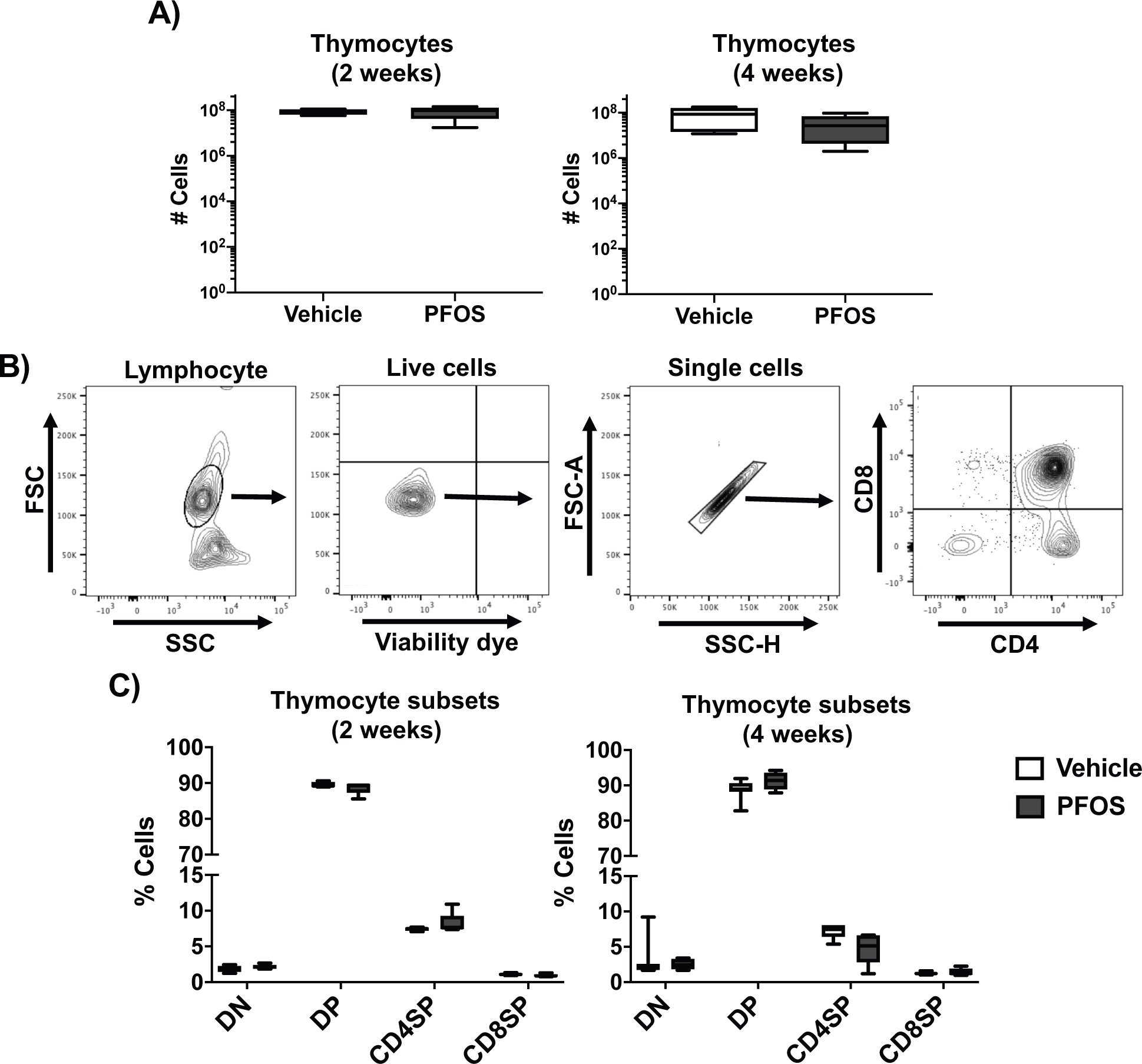
PFOS exposure at 3 μg/kg/day PFOS for 2 weeks or 1.5 μg/kg/day for 4 weeks does not significantly affect T cell development in the thymus. (**A**) Thymi from WT C57Bl/6 mice exposed to vehicle or PFOS at 3 μg/kg/day for 2 weeks (left panel) or 1.5 μg/kg/day for 4 weeks (right panel) were collected and thymocytes enumerated. Mean ± SEM, n=5 (vehicle exposed) and n=6 (PFOS exposed) for 2 weeks of exposure and n=7 (vehicle exposed) and n=5 (PFOS exposed). (**B**) Example flow cytometric plots of thymocytes and gating strategy to identify thymocyte populations. (**C**) CD4/CD8 double negative (DN), CD4/CD8 double positive, CD4 single positive (SP) and CD8SP populations plotted. Mean ± SEM, n=5 (vehicle exposed) and n=6 (PFOS exposed) for 2 weeks of exposure and n=7 (vehicle exposed) and n=5 (PFOS exposed).

### Effect of 2-week or 4-week exposure to PFOS on T and B cell development in the spleen

Once T lymphocytes develop in the thymus, they exit and migrate to peripheral organs, including the spleen and lymph nodes. We next analyzed T lymphocyte development in the spleen of the mice exposed to 3 μg/kg/day PFOS for 2 weeks or 1.5 μg/kg/day for 4 weeks for T cell subsets as indicated in figure 3A. We found no effect of PFOS at this concentration on total splenocytes (**Fig. 3A**), nor was there an effect on the percentage of αβ T cells, γδ T cells, invariant Natural killer T cells (*i*NKT cells), NK cells, and B cells (**Fig. 3B**, as analyzed by flow cytometric analysis exemplified in **Fig. 3C**). There was also no difference in the percentage of CD4^+^ and CD8^+^ αβ T cells (**Fig. 3D**) after 2 and 4 weeks of exposure. However, after 2 weeks of PFOS exposure, we observed a reduction in the percentage of naïve CD8^+^ αβ T cells, with an accompanying increase in memory CD8^+^ αβ T cells in the spleen, which was not observed in the CD4^+^ T cell compartment, or in either population at 4 weeks of exposure to PFOS (**Fig. 3D**, **E**). In addition, after 2 weeks of PFOS exposure, but not at 4 weeks, we observed an increase in the percentage of NK cells but not in the total number of NK cells (**Fig. 3D**). Analysis of B cell development for immature, mature and marginal zone B cells in the spleen revealed a reduction in the percentage of mature B cells in spleens from both 2 week and 4 week PFOS exposed animals, although the numbers of these cells were not affected (**Fig. 3F**).

**Figure 3.**
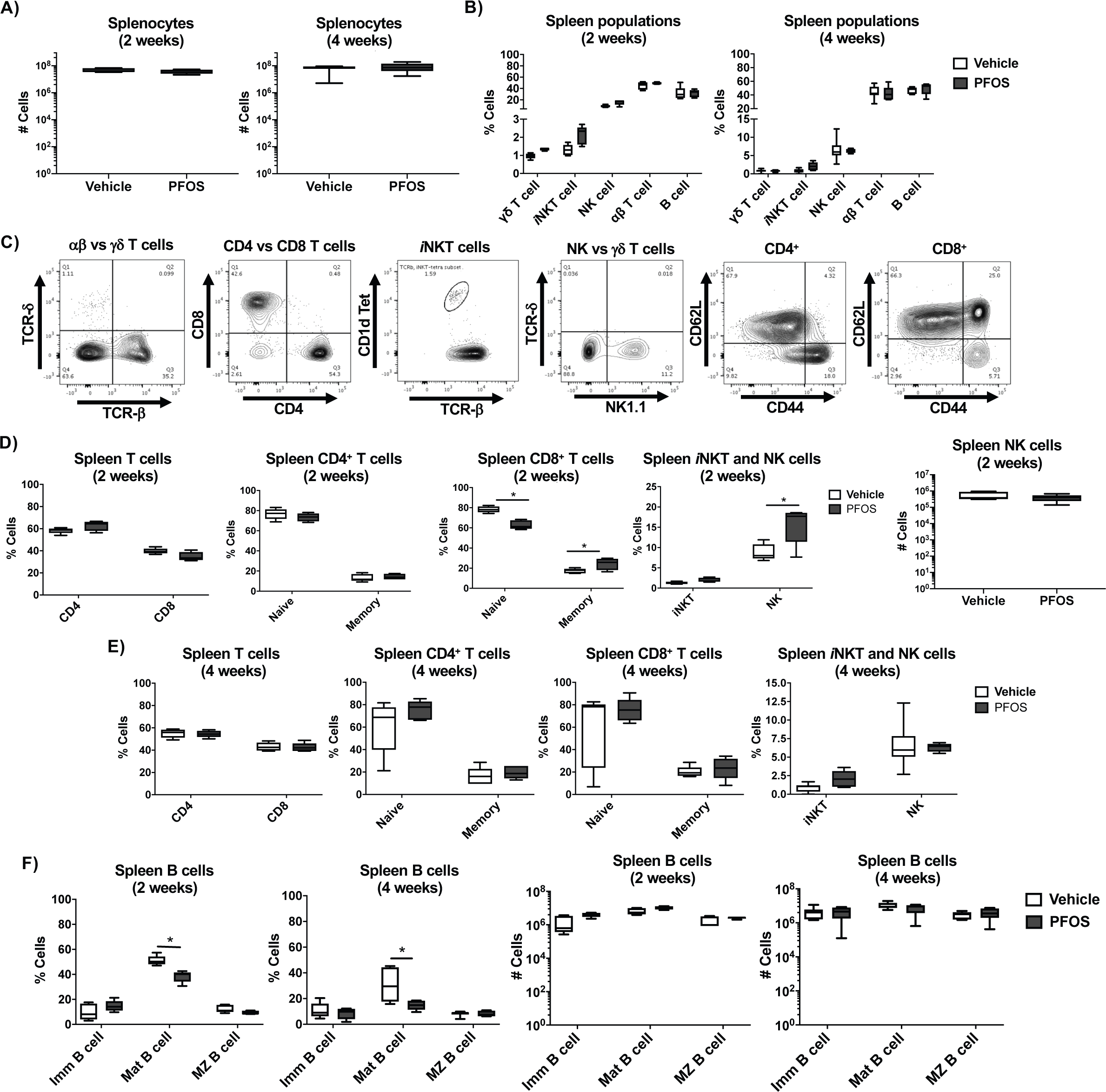
Effect of PFOS exposure at 3 μg/kg/day PFOS for 2 weeks or 1.5 μg/kg/day for 4 weeks on splenocyte populations. (**A**) Splenocytes from WT C57Bl/6 mice exposed to vehicle or PFOS at 3 μg/kg/day for 2 weeks (left panel) or 1.5 μg/kg/day for 4 weeks (right panel) were collected and enumerated. Mean ± SEM, n=5 (vehicle exposed) and n=5 (PFOS exposed) for 2 weeks of exposure and n=7 (vehicle exposed) and n=5 (PFOS exposed). (**B**) Analysis of the percentages of subpopulations of lymphocytes in spleens from mice described in (A). Mean ± SEM, n=5 (vehicle exposed) and n=5 (PFOS exposed) for 2 weeks of exposure and n=7 (vehicle exposed) and n=5 (PFOS exposed). (**C**) Example flow cytometric plots to identify T cell populations (*γδ*, *αβ*, CD4^+^, CD8^+^, *i*NKT, NK, and CD4^+^ and CD8^+^ naïve (CD44^lo^CD62L^hi^) and memory (CD44^Hi^) populations. (**D**) Analysis of the percentages of (left to right) CD4^+^ and CD8^+^ T cells (panel 1), naïve and memory subsets of CD4^+^ and CD8^+^ (panels 2 & 3) and *i*NKT and NK cells (panel 4), and number of NK cells (panel 5) in spleens from 2 week exposed mice described in (A). Mean ± SEM, n=5 (vehicle exposed) and n=5 (PFOS exposed), *p<0.0001 naïve CD8^+^ cells at 2 weeks, *p=0.0368 for memory CD8^+^ cells at 2 weeks vehicle vs PFOS exposed mice, *p=0.0014 for NK cells at 2 weeks, vehicle vs PFOS exposed mice. (**E**) Analysis of the percentages of (left to right) CD4^+^ and CD8^+^ T cells (panel 1), naïve and memory subsets of CD4^+^ and CD8^+^ (panels 2 & 3) and *i*NKT and NK cells (panel 4) in spleens from 4 week exposed mice described in (A). Mean ± SEM, n=5 (vehicle exposed) and n=5 (PFOS exposed). (**F**) Analysis of the percentages of (left to right) splenic immature, mature and marginal zone B cells after 2 weeks (panel 1) or 4 weeks of exposure (panel 2). Number of splenic immature, mature and marginal zone B cells after 2 weeks (panel 3) or 4 weeks of exposure (panel 4). Mean ± SEM, n=5 (vehicle exposed) and n=5 (PFOS exposed), *p=0.0003 for percent of mature B cells at 2 weeks, and *p=0.0014 for percent of mature B cells at 4 weeks vehicle vs PFOS exposed mice.

### Effect of 2-week or 4-week exposure to PFOS on T and B cell development in the bone marrow

As discussed above, hematopoietic stem cells differentiate into lymphoid progenitors, which develop into T and B cells. T cells develop in the thymus, and B cells develop in the bone marrow. In addition, some subsets of T cells, largely memory derived can also be found in the bone marrow. To further explore B and T cell development in the bone marrow, we analyzed cells collected from the bone marrow of mice exposed to 3 μg/kg/day PFOS for 2 weeks or 1.5 μg/kg/day for 4 weeks for B and T cell subsets. We found no effect of PFOS exposure for 2 or 4 weeks at this concentration on total leukocyte numbers in the bone marrow (**Fig. 4A, D**). There was also no effect on the number of granulocytes and lymphocytes, including T and B cells after 2 weeks of exposure (**Fig. 4B**). After 2 weeks of exposure there was no effect on the proportion of B cells, and CD4^+^ and CD8^+^ subpopulations of T cells (**Fig. 4C**), nor on naïve and memory CD4^+^ T cells, although there was an increase in the proportion of CD8^+^ naïve T cells, and a decrease in the proportion CD8^+^ memory T cells (**Fig. 4C**). We also found that after 4 weeks of exposure there was a decrease in the proportion of CD4^+^ T cells and an increase in CD8^+^ memory T cells (**Fig. 4F**), as well as an increase in proportion of the NK cells at 4 weeks, although we found no difference in the number of NK cells (**Fig. 4G**).

**Figure 4.**
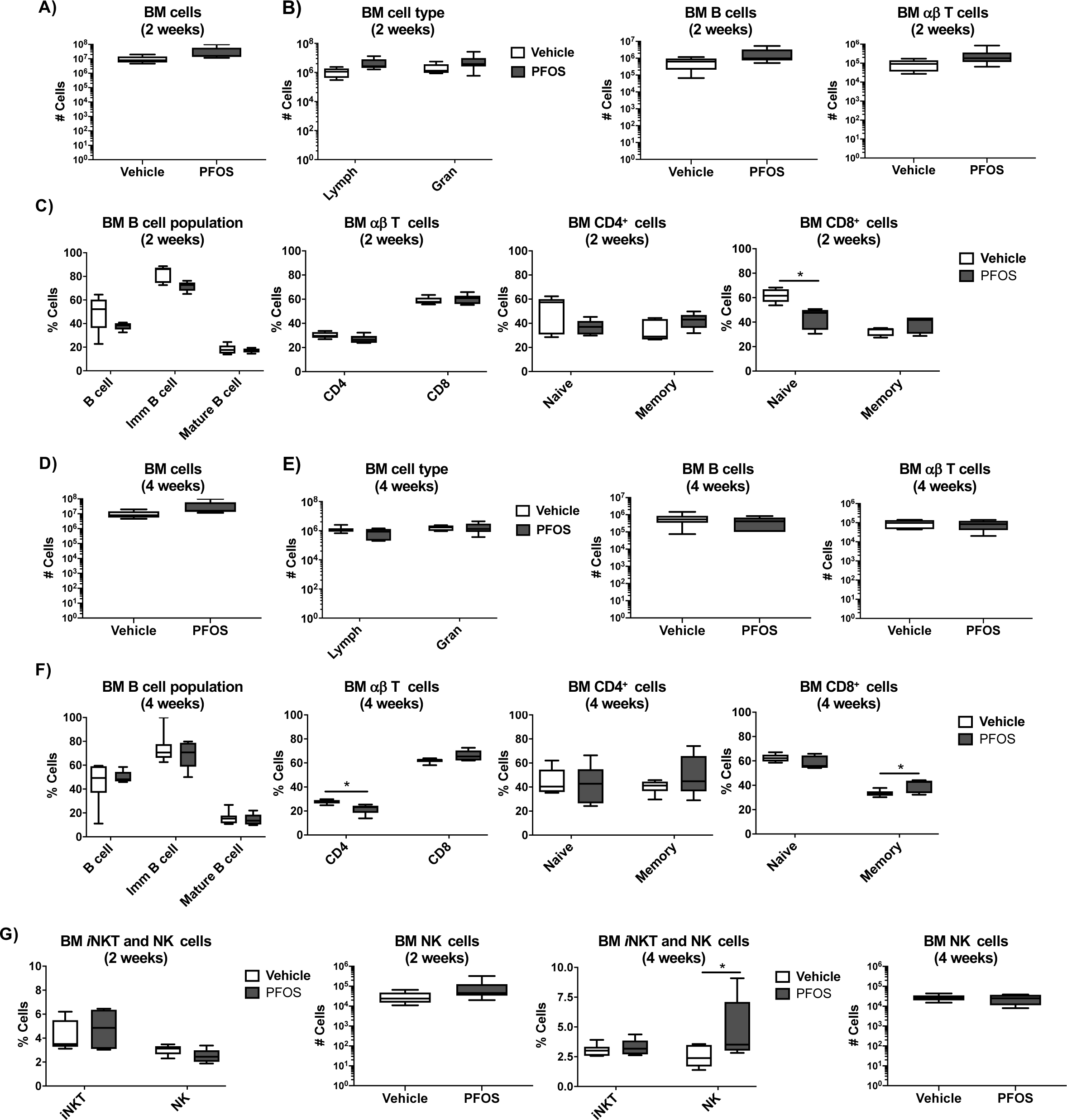
Effect of PFOS exposure at 3 μg/kg/day PFOS for 2 weeks or 1.5 μg/kg/day for 4 weeks on bone marrow lymphocyte populations. (**A, D**) Bone marrow lymphocytes from WT C57Bl/6 mice exposed to vehicle or PFOS at 3 μg/kg/day for 2 weeks (**A**) or 1.5 μg/kg/day for 4 weeks (**D**) were collected and enumerated. Mean ± SEM, n=5 (vehicle exposed) and n=5 (PFOS exposed) for 2 weeks of exposure and n=7 (vehicle exposed) and n=5 (PFOS exposed) for 4 weeks. (**B**) Analysis of the percentages of subpopulations of lymphocytes in spleens from mice described in (A). (**C**) Analysis of the percentages of (left to right) B cells (panel 1), *αβ* T cells (panel 2), CD4^+^ and CD8^+^ naïve and memory subsets (panels 3&4) in spleens from 2 week exposed mice described in (A). *p=0.0005 for naïve CD8^+^ cells at 2 weeks vehicle vs PFOS exposed mice. (**E**) Analysis of the percentages of subpopulations of lymphocytes in spleens from mice described in (D). (**F**) Analysis of the percentages of (left to right) B cells (panel 1), *αβ* T cells (panel 2), CD4^+^ and CD8^+^ naïve and memory subsets (panels 3&4) in spleens from 4 week exposed mice described in (D). *p=0.0107 for CD4^+^ *αβ* T cells and *p=0.0382 for memory CD8^+^ cells for vehicle vs PFOS exposed mice. (**G**) Analysis of the percentages (panels 1&3) and number (panel 2&4) of *i*NKT and NK cells in spleens from 2 week exposed mice described in (A), (panels 1&2), or 4 week exposed mice described in (D), (panels 3&4). *p=0.0149 for NK from 2 week exposed vehicle vs PFOS exposed mice.

### Effect of 2-week or 4-week exposure to PFOS on T cells in the liver

The liver has been shown to be a target for PFOS at higher concentrations, and while we did not observe changes in liver weights at this concentration, we examined T cells in this organ. We found that mice exposed to PFOS for 2 but not 4 weeks had higher numbers of total liver lymphocytes (**Fig. 5A, D**), although there was no difference in number or percentages of liver T cells, *i*NKT cells or NK cells (**Fig. 5B, E**). There was also no difference in percentages of CD4^+^ and CD8^+^ subpopulations of T cells (**Fig. 5C, F**), *i*NKT cells, NK cells, nor on naïve and memory CD4^+^ T cells, with the exception of a decrease in percentages of CD8^+^ naïve T cells after 2 but not 4 weeks of exposure (**Fig. 5C, F**).

**Figure 5.**
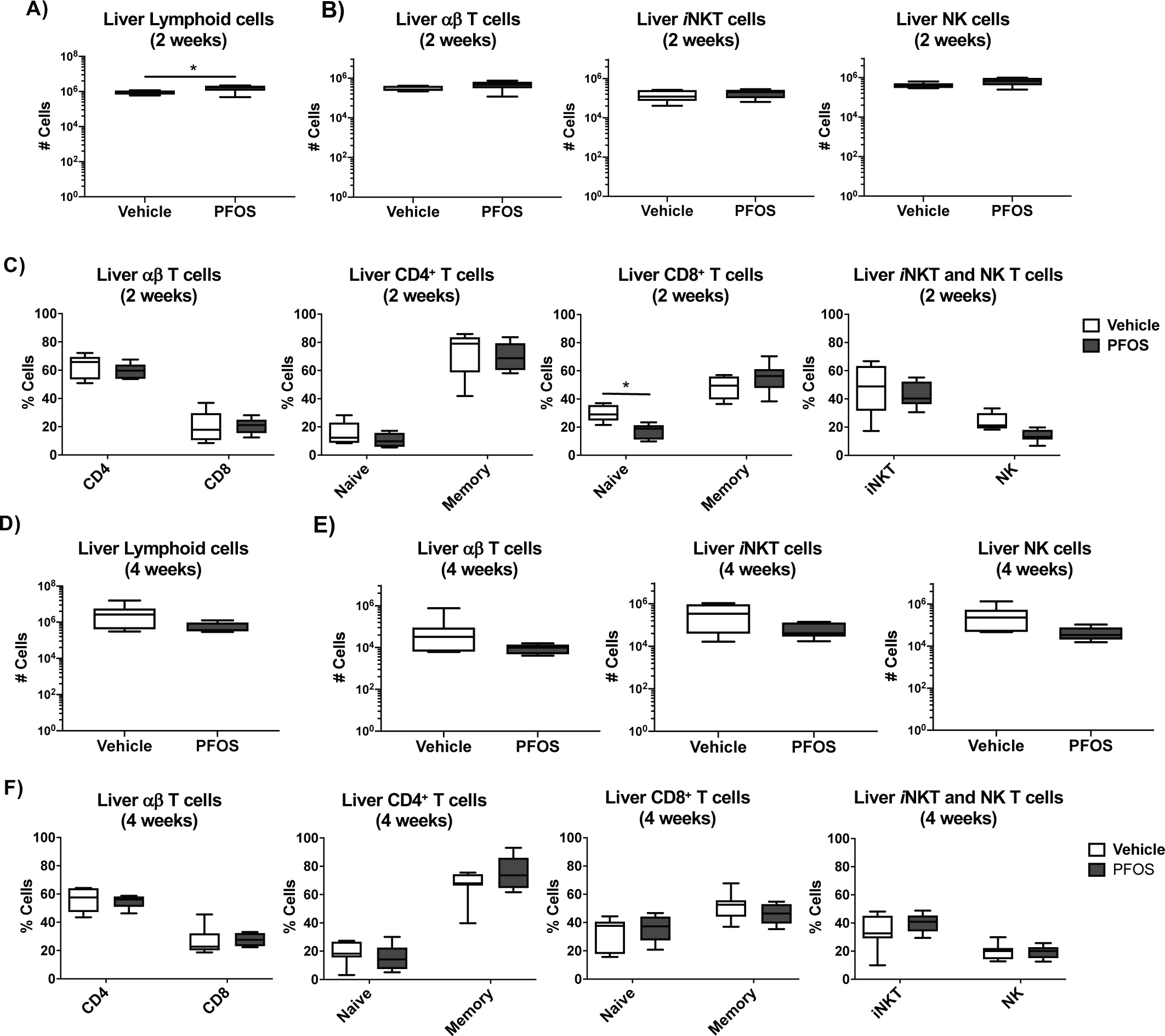
Effect of PFOS exposure at 3 μg/kg/day PFOS for 2 weeks or 1.5 μg/kg/day for 4 weeks on liver lymphocyte populations. (**A, D**) Number (Mean ± SEM) of liver lymphocytes from WT C57Bl/6 mice exposed to vehicle or PFOS at 3 μg/kg/day for 2 weeks (n=5 (vehicle exposed) and n=6 (PFOS exposed)) (**A**, p=0.0450), or 1.5 μg/kg/day for 4 weeks (n=7 (vehicle exposed) and n=5 (PFOS exposed)) (**D**). (**B, E**) Number (Mean ± SEM) of liver *αβ* T cells (left panel), *i*NKT cells (middle panel) or NK cells (right panel). (**C, F**) Percentage of CD4^+^, CD8^+^ (left panel), CD4^+^ and CD8^+^ naïve and memory cells (middle panels, p=0.0320 for naive CD8^+^ cells after 2 week exposure), and *i*NKT and NK cells (right panel).

### 4-week exposure to 1.5 μg/kg/day PFOS does not affect viral clearance and the immune response to influenza virus infection

The immune response to influenza virus is critical for protection against infection. We therefore determined the effect of PFOS exposure on the response to influenza virus infection. 8-week-old C57BL/6 male and female mice were exposed to vehicle or 1.5 μg/kg/day PFOS for 4 weeks. Twenty four hours after the last exposure, animals were infected with influenza virus (Influenza A/WSN/33(H1N1) (**Fig. 6A-D**), or a mixture of influenza A/WSN/33(H1N1) (WSN-OVA_I_) and influenza A/WSN/33(H1N1) (WSN-OVA_II_), or PBS as control (**Fig. 6E-J**). In the first experiment where mice were infected with Influenza A/WSN/33(H1N1) (**Fig. 6A-D**), PFOS exposure did not significantly affect virus induced weight loss (**Fig. 6A**). Similarly, there was no effect of PFOS exposure at this concentration on the amount of virus present in the lung upon sacrificing at 11 days post infection (**Fig. 6B**). There was also no difference in the number of inflammatory cells, T cells and granulocytes, in the lung (**Fig. 6C**), nor the proportion of these cells in the BALF or lung (**Fig. 6D**). Similarly, there was no difference in the proportion of CD4^+^ or CD8^+^ T cells, nor CD4^+^CD44^Hi^, CD8^+^CD44^Hi^ (with CD44^Hi^ cells representing activated or memory cells) in BALF (**Fig. 6D**, top panels), although PFOS exposed mice exhibited reduced proportions of CD4^+^ T cells and increased CD8^+^ T cells, with increased proportion of CD4^+^CD44^Hi^ in the lung (**Fig. 6D**, bottom panels). In addition, there was no difference in the proportion of TNF*α*^+^ CD4^+^ or CD8^+^ T cells in the lung (as analyzed by intracellular cytokine staining following PMA/Ionomycin stimulation in vitro) (**Fig. 6D**).

**Figure 6.**
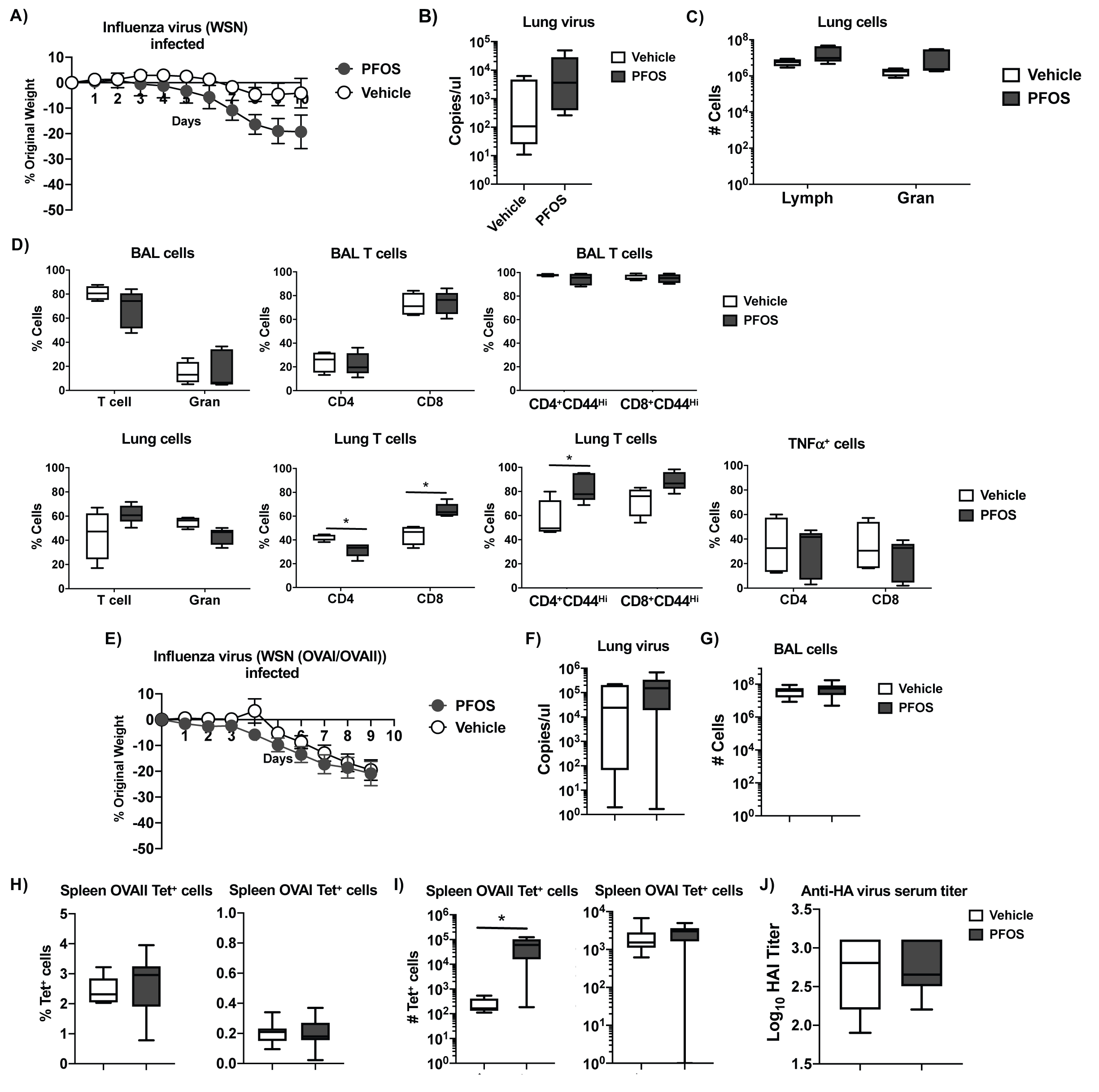
Effect of PFOS exposure at 1.5 μg/kg/day for 4 weeks on response to influenza infection. (**A, D**) WT C57Bl/6 mice exposed to vehicle or PFOS at 1.5 μg/kg/day were infected with influenza WSN (**A**), or mixture of WSN-OVA_I_ and WSN-OVA_II_ (**E**) on day 0, and weight loss as a percentage of original weight determined over the indicated days. WSN infected n=4 (vehicle exposed) and n=5 (PFOS exposed), WSN-OVA_I_ and WSN-OVA_II_ n=7-10. (**B, F**) Analysis of influenza virus at day 11 (WSN, (B)) or day 10 (WSN-OVA_I_ and WSN-OVA_II_, (F)) by qRT-PCR of viral genome. (**C**) Number of lung lymphocytes and granulocytes. (**D**) Percentage of T cells and granulocytes, CD4^+^ and CD8^+^ T cells, and CD44^Hi^ CD4^+^ and CD8^+^ T cells in BAL (top panels) and lung (bottom panels). Percentage of lung TNF*α*^+^ CD4^+^ and CD8^+^ T cells T cells (bottom left panel). *p=0.0346 for percent lung CD4^+^ T cells, *p=0003 for percent lung CD8^+^ T cells, *p=0.0110 for CD44^Hi^ CD4^+^ T cells. (**G**) Number of BAL cells following infection with WSN-OVA_I_ and WSN-OVA_II_. (**H**) Percentage of OVA/MHCII tetramer specific CD4s^+^ (left panel) and OVA/MHCI tetramer specific CD8^+^ T cells (right panel) in spleen. (**I**) Number of OVA/MHCII tetramer specific CD4^+^ (left panel) and OVA/MHCI tetramer specific CD8^+^ T cells (right panel) in spleen. (**J**) HAI titers of serum from mice used in figure 6E.

In a second experiment mice were infected with a mixture of influenza A/WSN/33(H1N1) (WSN-OVA_I_) and influenza A/WSN/33(H1N1) (WSN-OVA_II_) so as to follow the antigen specific CD8^+^ and CD4^+^ T cell response using MHC-class I/OVA and class II/OVA tetramers that identify OVA-specific CD4^+^ and CD8^+^ T cells (**Fig. 6E-J**). As in the experiment where mice were infected with influenza A/WSN/33(H1N1) (see **Fig. 6A**), infection with WSN-OVA_I_ and WSN-OVA_II_ following PFOS exposure did not significantly affect virus induced weight loss (**Fig. 6E**). Similarly, there was no effect of PFOS exposure at this concentration on the amount of virus present in the lung upon sacrificing at 11 days post infection (**Fig. 6F**). There was also no difference in the number of inflammatory cells in the BALF collected from flu infected mice regardless of exposure to PFOS (**Fig. 6G**). Analysis of the antigen specific T cell response using the MHC-class I/OVA and class II/OVA tetramers revealed no difference in the percentage of antigen specific CD4^+^ or CD8^+^ T cells in the spleens of these mice, although PFOS exposed mice exhibited a higher number of antigen specific CD4^+^ T cells (**Fig. 6H, I**). Analysis of serum for neutralizing antibodies generated against the virus by Hemagglutinin Agglutination Inhibition (HAI) assay revealed no difference in anti-viral titers between vehicle and PFOS exposed mice at this concentration (**Fig. 6J**).

## Discussion

The persistent nature of PFOS has led to the investigation of possible health effects of exposure in humans and laboratory animals. Here we have explored the effects of PFOS exposure (3 and 1.5 μg/kg/day of PFOS for 2 and 4 weeks, respectively (equivalent to ∼100 ng/ml serum PFOS)) on immune cell development and function in mice. We find that at this concentration, oral exposure of mice to PFOS did not result in weight loss during the 2 and 4 week exposure. We also find that PFOS does not affect T-cell development during this time. However, we find that PFOS exposure reduced immune cell populations in some organs, and increased their numbers in others. We find that exposure of mice to PFOS at 1.5 μg/kg/day of PFOS for 4 weeks does not affect weight loss or survival following infection with influenza virus, nor does it affect viral clearance, nor the antigen specific antibody and T cell response. These results suggest that exposure of mice to PFOS at 3 and 1.5 μg/kg/day of PFOS for 2 and 4 weeks does not suppress the immune system or its response to viral infection.

Based on reports of PFOS-mediated suppression of the antibody response in animal studies, and epidemiological studies in humans, the NTP previously concluded that PFOS is “presumed to be an immune hazard to humans”^12^. Previous studies using exposure to higher levels of PFOS have suggested an effect on the immune system, with either no effect on immune cell numbers, reduced thymic and splenic T (including reduced CD8^+^ and CD4^+^ T cells) and B cells, reduced or no effect on T cell proliferation, reduced IL2 and IFN*γ*, (Th1 cytokines) increased IL4 (Th2 cytokine), decreased TNF*α*, no effect on IL10, increased or no effect on IL1*β*, increased or decreased IL6, increased or decreased NK cell activity (depending on exposure concentration), increased serum IgG, or reduced IgM, but increased IgG, IgG_1_, and IgE levels in response to SRBC injection^24–34^. In many of these studies, animals were in general exposed to PFOS concentrations >5 mg/kg/day and in some cases >50 mg/kg/day. At high levels of PFOS exposure (e.g. >20 mg/kg/day for 7 days, equivalent to >280,000 ng/ml) and in a dose-dependent manner, mice experienced decreased food intake, increased liver mass, weight loss and serum corticosteroid; the latter could affect immune populations^33, 35^. At concentrations in the range used in this study, little to no effect on liver weights and body weights, or on the immune system has been reported^30, 36^. Here, following exposure to 2 week or 4 weeks leading to serum concentrations of ∼100-125 ng/ml (for a TAD of 0.14 mg/kg after 4 weeks), with no weight loss or changes in organ weights, we found no effect on T cell development in the thymus, including on the percentages of double negative, double positive or CD4 and CD8 single positive thymocytes.

Analysis of spleen also revealed no effect on total cell numbers, however after 2 weeks, but not 4 weeks of PFOS exposure, we observed a reduction in the percentage of naïve CD8^+^ αβ T cells, with an accompanying increase in memory CD8^+^ αβ T cells, which was not observed in the CD4^+^ T cell compartment. This transient change could be due to changes in these populations, or changes in the recirculation patterns of these T cell subsets. Indeed, we also observed an increase in the CD8^+^ naïve T cells, and a decrease in CD8^+^ memory T cells in the bone marrow after 2 weeks of PFOS exposure. In addition, after 2 weeks of PFOS exposure, but not after 4 weeks, we observed an increase in the percentage of NK cells but not in the total number of NK cells. Analysis of B cells in the spleen revealed a reduction in percentage of mature B cells in the spleens from both 2 week and 4 week PFOS exposed animals, although the number of these cells were not affected. It is possible that accompanying changes in other B cell subsets also occur that affect the proportions of these B cell subsets. The observed changes may also reflect changes in recirculation patterns of different cell subsets that may occur during normal immune homeostasis.

Our analysis of the effect of PFOS on bone marrow populations of T cells revealed a decrease in proportion of CD4^+^ T cells and an increase in CD8^+^ memory T cells after 4 weeks of PFOS exposure, as well as an increase in the proportion of NK cells, although again, no difference in the number of NK cells. No effect was observed on immature and mature B cells. Studies of exposure of mice to oral PFOS (in food) have revealed dose-dependent increase in PFOS multiple tissues including blood, liver and bone marrow, with liver retaining the highest levels^37^. In the liver, we found an increase in the number of lymphocytes at 2 weeks, but not at 4 weeks, with no difference in the numbers or percentages of liver T cells, *i*NKT cells or NK cells. We also found a transient decrease in naïve CD8^+^ T cells after 2 weeks, but not after 4 weeks of exposure. Other researchers have examined the effects of PFOS on bone marrow cells (at levels of 0.002% w/w PFOS for 10 days, equivalent to 3.1 mg/kg) and have also reported no effect on cell numbers or B cell development^38^. Our work has also found no effect of PFOS exposure on the numbers or percentages of these cells at the levels of PFOS exposure used in these experiments.

Little work has been done exploring the effects of PFOS on response to infection in mice. Suo et al, exposed mice to much higher levels of PFOS (2 mg/kg/day) and examined the response to *Citrobacter rodentium* (CR) infection, and found enhanced early response to CR, but eventual delayed clearance of bacteria^39^. However, these high levels of PFOS exposure may have other effects as discussed previously. Guruge et al reported that doses in a similar range to what was used in this study at 5 (∼189 ng/ml serum levels) and 25 mg/kg/day (∼670 ng/ml serum levels) PFOS exposure was accompanied by increased weight loss and mortality following infection with the mouse adapted influenza A virus (100 pfu A/PR/8/34 (H1N1) strain)^40^. However, there was no analysis of immune response in these experiments^40^.

Here, we have also explored the effect of PFOS exposure on the response to influenza infection, using the mouse adapted A/WSN/33 (H1N1) strain of influenza. We found no significant effect on infection-induced weight loss, virus levels in the lung, number of lung T cells and granulocytes. We did find that PFOS exposed mice exhibited reduced proportions of lung CD4^+^ T cells and increased CD8^+^ T cells, although there was an increased proportion of activated/memory CD4^+^ T cells (as determined by higher levels of CD44). In addition, there was no difference in ex vivo stimulated production TNF*α* by T cells in the lung. We also analyzed T cell specific antigen responses to flu infection by using recombinant influenza viruses A/WSN/33 (H1N1) (WSN-OVA_I_) and influenza A/WSN/33(H1N1) (WSN-OVA_II_). We found that PFOS exposure did not significantly affect the percentage of antigen specific CD4^+^ or CD8^+^ T cells in the spleens of infected mice, although PFOS exposed mice exhibited a higher number of antigen specific CD4^+^ T cells. Finally, our analysis of serum from infected for virus specific antibodies, revealed no effect of PFOS exposure. Our results suggest that at this concentration of PFOS, there is no suppression of the antiviral response to flu infection. It is possible that our results differ due to the different strains of mice used (our studies used C57Bl/6, the previous study used B6C3F1), as different strains of mice exhibit differential susceptibility to flu infection^41, 42^. The viral strain may also be a potential explanation as our study used A/WSN/33 (H1N1) and the previous study used A/PR/8/34 (H1N1)), although they are reported to have similar LD_50_s in mice^41^.

## Conclusion

In this study, we have examined the effects of subchronic exposure of PFOS on immune cell development and function in mice. At approximately 100-125 ng/mL of serum PFOS, it has minimal effects on lymphocyte subpopulation proportions or numbers, and does not suppress the immune response to influenza infection. Further studies in laboratory animals, as well as in humans, need to be considered in order to fully determine the potential effects of PFOS on immune system development and response.

## Acknowledgement

We thank members in the August lab for helpful discussions. This work was supported by an unrestricted university research grant from 3M Company (to A.A.). B.I is supported by the Cornell IMSD program (R25GM125597 to A.A.). C.L. is a Cornell Sloan Scholar.

## Author contributions

L.T., A.R., C.L., S.C. and B.I. conducted experiments; L.T., B.I, S.C. and A.A. designed research, L.T., B.I, S.C. and A.A. analyzed data, and L.T., S.C. and A.A. wrote the manuscript.

